# Improvement in Neoantigen Prediction via Integration of RNA Sequencing Data for Variant Calling

**DOI:** 10.1101/2023.07.02.547404

**Authors:** Bui Que Tran Nguyen, Thi Phuong Diem Tran, Huu Thinh Nguyen, Thanh Nhan Nguyen, Thi Mong Quynh Pham, Hoang Thien Phuc Nguyen, Duc Huy Tran, Vy Nguyen, Thanh Sang Tran, Truong-Vinh Ngoc Pham, Minh-Triet Le, Minh-Duy Phan, Hoa Giang, Hoai-Nghia Nguyen, Le Son Tran

## Abstract

Neoantigen-based immunotherapy has emerged as a promising strategy for improving the life expectancy of cancer patients. This therapeutic approach heavily relies on accurate identification of cancer mutations using DNA sequencing (DNAseq) data. However, current workflows tend to provide a large number of neoantigen candidates, of which only a limited number elicit efficient and immunogenic T-cell responses suitable for downstream clinical evaluation. To overcome this limitation and increase the number of high-quality immunogenic neoantigens, we propose integrating RNA sequencing (RNAseq) data into the mutation identification step in the neoantigen prediction workflow. In this study, we characterize the mutation profiles identified from DNAseq and/or RNAseq data in tumor tissues of 25 patients with colorectal cancer (CRC). We detected only 22.4% of variants shared between the two methods. In contrast, RNAseq-derived variants displayed unique features of affinity and immunogenicity. We further established that neoantigen candidates identified by RNAseq data significantly increased the number of highly immunogenic neoantigens (confirmed by ELISpot) that would otherwise be overlooked if relying solely on DNAseq data. In conclusion, this integrative approach holds great potential for improving the selection of neoantigens for personalized cancer immunotherapy, ultimately leading to enhanced treatment outcomes and improved survival rates for cancer patients.

## INTRODUCTION

Colorectal cancer (CRC) is a major global health concern, being the third most common cancer in the world and the fifth leading cause of cancer-related mortality among the Vietnamese population (1, 2). Traditional treatments, such as surgery, chemotherapy, and radiation therapy, have limited efficacy and are poorly tolerant, particularly in advanced stages of CRC (3). Immunotherapy, while not a cure for CRC, has the potential to significantly improve patient survival rates and quality of life (4, 5). In metastatic CRC patients, immunotherapy has demonstrated promise in improving outcomes. Immune checkpoint inhibitors (ICIs), which block negative regulatory pathways in T-cell activation, have been approved by the US Food and Drug Administration (FDA) for the treatment of deficient mismatch repair (dMMR) or high microsatellite instability (MSI-H) CRC patients (6–8). However, alternative immunotherapy strategies are urgently required for CRC patients, as patients with proficient mismatch repair (pMMR) or microsatellite stability (MSS) have not shown significant responses to immune checkpoint inhibitors (6, 9).

Neoantigens (neopeptides) have emerged as potential targets for personalized cancer immunotherapy, including CRC (10, 11). Neoantigens are peptides that result from somatic mutations and can be displayed by class I human leukocyte antigen (HLA-I) molecules on the surface of cancer cells, thereby activating immune-mediated tumor killing (12). Recent studies have demonstrated that the presence of neoantigens is associated with better responses to immune checkpoint inhibitor (ICI) therapy in CRC patients (13, 14). A high neoantigen burden has been linked to improved overall survival and progression-free survival in patients with various solid tumors, including CRC (13, 14). Therefore, neoantigen-based immunotherapies are thought to have the potential to significantly improve treatment outcomes for CRC patients.

The identification of neoantigens with strong binding affinity to their respective HLA-I molecules and high immunogenicity is critical for the development of effective neoantigen-based therapies. This process involves the use of next-generation sequencing (NGS) and bioinformatics tools. Initially, DNA sequencing of tumor tissues and paired white blood cells enables the identification of cancer associated genomic mutations, while RNA sequencing is used to determine patient’s HLA-I allele profile and to quantify expression levels of genes carrying mutations. Next, tumor somatic variant, HLA-I allele, and gene expression data are analyzed using *in silico* tools based on machine learning algorithms to predict the binding affinity of neoantigens to patients’ HLA-I alleles and their potential to activate T cell responses (15–17). This standard workflow has been exploited in numerous studies to identify clinically relevant neoantigens in melanoma, lung cancer, and other malignancies (16, 18).

Despite promising results, only small portions of patients benefit from the current approach due to the limited number of effective immunogenic neoantigens identified for each patient. To maximize the detection of potential neoantigens, whole exosome sequencing (WES) has been employed to comprehensively profile the cancer-specific landscape (19–21). While WES allows a much larger search space for mutations within the genome, it is not a cost- and time-effective approach. Moreover, a significant proportion of identified tumor DNA mutations, especially those which are not actively transcribed or transcribed at very low levels, might not result in the formation of neoantigens (18). Lastly, WES-based mutation calling is inefficient in capturing all tumor somatic mutations, especially clonal mutations with low frequencies and underrepresentation in the sequencing data (22), while targeting combined neoantigens derived from both clonal and subclonal mutations is necessary to evoke efficient immune-mediated cell death in a broader range of tumor cells. Therefore, relying solely on DNAseq data for tumor mutation calling, which has traditionally been the basis for identifying neoantigens, may not capture the full extent of tumor-related mutations, resulting in an incomplete identification of neoantigens.

Genetic variants at the RNA level are frequently excluded from conventional bioinformatic workflows, despite several studies indicating that neoantigens can be derived from RNA mutations, such as splicing, polyadenylation dysregulation, or RNA editing (23, 24). In addition, recent studies have shown that the presence of variant-bearing transcripts is an important factor for accurate identification of immunogenic neoantigen candidates (25, 26). Therefore, integrating RNAseq data into tumor mutation calling holds promise for unveiling a more comprehensive repertoire of neoantigens and, consequently, advancing the development of personalized immunotherapies for cancer. However, the feasibility and effectiveness of this approach require further examination.

To assess the utility of RNAseq analysis for neoantigen identification, we compared the cancer mutation profiles, binding affinity to HLA-I of neoantigens identified from RNAseq and DNAseq, and their predicted immunogenicity across 25 CRC patients. Moreover, we performed experimental validation to assess the effectiveness of utilizing RNAseq for the identification of immunogenic neoantigens. This validation utilized the ELISpot assay to measure the ability of neoantigen candidates, predicted from DNAseq and RNAseq-derived variants, to activate T cells in PBMCs obtained from four CRC patients.

## MATERIALS AND METHODS

### Tumor biopsy and peripheral blood collection

A total of 25 patients diagnosed with colorectal cancer (CRC) were enrolled in this study from the University Medical Center at Ho Chi Minh city between June 2022 and April 2023. The confirmation of CRC was based on abnormal mammograms and histopathological analysis confirming the presence of malignancy. The stages of CRC were determined following the guidelines provided by the American Joint Committee on Cancer and the International Union for Cancer Control. Prior to participation, all patients provided written informed consent for the collection of tumor and whole blood samples. Relevant clinical data, including demographics, cancer stages, and pathology information, were extracted from the medical records of the University Medical Center. Detailed information regarding the clinical factors of the patients can be found in **Table S1**. The Ethic Committee of University of Medicine and Pharmacy at Ho Chi Minh City, Vietnam approved this study. Fresh CRC specimens were collected immediately after biopsy or tumor resection and were placed in microtubes containing RNAlater, an RNA stabilization solution (Thermo Fisher Scientific, Japan). For four patients, ten mL of peripheral blood were collected serially before surgery and stored in Heparin tubes.

### Targeted DNA and RNA sequencing

The DNA/RNA samples were isolated using either the AllPrep DNA/RNA Mini Kit or the AllPrep DNA/RNA/miRNA Universal Kit (Qiagen, Germany) as per the manufacturer’s protocol. In addition, matched genomic DNA from the white blood cells (WBC) of individual was also extracted from the buffy coat using the GeneJET Whole Blood Genomic DNA Purification Mini kit (ThermoFisher, MA, USA), following the manufacturer’s instructions. Genomic DNA samples from the patients’s paired tumor tissues and WBCs were used to prepare DNA libraries for DNA sequencing with the ThruPLEX Tag-seq Kit (Takara Bio, USA). The libraries were then pooled and hybridized with pre-designed probes for 95 targeted genes (Integrated DNA Technologies, USA). This gene panel encompasses commonly mutated genes in CRC tumors, as reported in the Catalog of Somatic Mutations in Cancer (COSMIC) database. The DNA libraries were then subjected to massive parallel sequencing on the DNBSEQ-G400 sequencer (MGI, Shenzhen, China) for paired-end reads of 2x100 bp with an average target coverage of 200X (with actual coverage from 89 to 968X).

Isolated total RNA was subjected to a NEBNext Poly(A) mRNA Magnetic Isolation Module (New England Biolabs, MA, USA) to isolate intact poly(A)+ RNA as per manufacturer instructions. RNA libraries were constructed using NEBNext Ultra Directional RNA Library Prep Kit for Illumina (New England Biolabs). These libraries were subsequently sequenced for paired end reads of 2x100 bp on an MGI system at 50X depth coverage.

### Variant calling from DNAseq and RNAseq data

To select the optimal variant calling tool for DNA-seq data, we evaluated the performance of three different pipelines including Dragen, VarScan and MuTect2, which are commonly used for somatic variant calling (27, 28). Among the three pipelines, Dragen demonstrated superior performance for detecting a set of validated ground truth variants in a standard dataset downloaded from a public repository, NCBI Sequencing Read Archive SRA (ID: SRR7890830) (**Figure S1A**). Therefore, we utilized Dragen (Illumina) (29) in tumor-normal mode to call somatic mutations from DNAseq data. The default filtering thresholds of Dragen were used to call SNPs and indels. SNPs were further filtered using the dbSNP and 1000 Genome datasets. Germline mutations in tumor tissues were identified by comparing them with matched WBC-DNA samples. Mutations within immunoglobulin and HLA genes were excluded due to alignment difficulties in these highly polymorphic regions that require specialized analysis tools (30). Additionally, synonymous mutations were removed from downstream analysis. Included for analysis were somatic mutations that surpassed a minimum threshold of ≥2% variant allele frequency (VAF) in DNA extracted from fresh frozen tissues.

To identify the most suitable variant calling tools for RNAseq data, we assessed the performance of two different pipelines, VarScan and MuTect2 by comparing the proportions of variants that overlapped with DNA-derived variants. Sequencing reads were trimmed using Trimmomatic (31) and aligned to the human reference genome using STAR (version 2.6.0c) (32). Prior to alignment, the raw sequencing reads underwent quality checks using FastQC version 0.11.9 (33). VarScan 2 (27), which accepts both DNA and RNA-seq data, was used to call mutations in paired tumor and WBC samples in 95 cancer-associated genes, again in the tumor-normal mode. Four filtering steps were applied: (i) only calls with a PASS status were used, (ii) population SNPs overlapping with a panel of normal samples from the 1000 Genome dataset were excluded, (iii) somatic mutations included for analysis met a minimum threshold of ≥10× read depth and ≥2% VAF in RNA extracted from FF tissue, and (iv) synonymous mutations and those related to HLA were removed from downstream analysis. The resulting BAM files were sorted and indexed using Samtools version 1.10 (34), and PCR duplicates were eliminated using Picard tools version 2.25.6 (35). The mutations from RNAseq data were also called using MuTect2, a variant caller from the Genome Analysis Toolkit (GATK) pipeline. Like VarScan, the MuTect2 pipeline was run in tumor versus normal mode, utilizing default settings. Following variant calling, a similar variant filtration step was also applied to eliminate potential false positives. Somatic variants from the two pipelines were manually checked using Integrative Genomics Viewer (v2.8.2). The VCF files generated by Dragen (for DNAseq) and by MuTect2 and VarScan (for RNAseq) were subsequently annotated using the Ensembl Variant Effect Predictor (VEP version 105) (36) to extract the potential effect of variants on the phenotypic outcome.

### Gene expression quantification and tumor purity estimation

We used the Cufflinks (37) to analyze the tumor RNAseq data using the Ensemble human reference transcriptomes (GRCh38) for assessing gene expression. The expression data was used to calculate the tumor purity via ESTIMATE (v1.0.13) package, (R-v3.6.3) (38).

### *In silico* prediction of HLA binding affinity and immunogenicity

Class I HLA alleles (HLA-A/B/C) with two-digit resolution were identified from patient tumor RNA-seq data using OptiType tool (39). The annotated VCF files were analyzed using pVAC-Seq, a tool of pVACtools (v1.5.9) (15, 40, 41) with the default settings, except for disabling the coverage and MAF filters. We used all HLA-I binding algorithms that were implemented in pVAC-Seq to predict 8 to 11-mer epitopes binding to HLA-I (A, B, or C) for downstream analysis. Neoantigens were subjected to MHC binding predictions and subsequent prioritization based on their binding affinity scores (measured in nM) using NetMHCpan-4.1 (17). The prioritization process involved calculating the percentile ranking of each neoantigen’s binding affinity score within the distribution of scores for the corresponding HLA allele. Neoantigens with a percentile rank lower than 2% were prioritized.

The immunogenicity of neoantigens was validated by the PRIME tool (42) with default settings. To predict the immunogenicity of neoantigens, a two-step ranking process was employed, involving ranking the neoantigens based on their immunogenicity score and estimating percentiles for each HLA allele. These scores represented the predicted likelihood of a neoantigen being immunogenic. The neoantigens were then ranked in descending order based on their immunogenicity scores, enabling the prioritization of neoantigens with higher predicted immunogenicity for further analysis. A ranking value for immunogenicity was assigned to each neoantigen candidate by determining the percentile rank of its immunogenicity score within the group of neoantigens predicted to bind to the same HLA allele. To identify public neoantigens, we conducted a comprehensive search of several databases, including TSNAdb (43, 44), NeoPeptide (45), dbPepNeo (46, 47), NEPdb (48), TANTIGEN (49, 50), and IEDB ((51). All databases contained epitopes from published studies where their immunogenicity was validated by immunological assays.

### Isolation, culture, and stimulation of PBMCs with long peptides

Peripheral blood samples from four patients were collected prior to surgery using BD Vacutainer Heparin Tubes (BD Biosciences, NJ, USA). Peripheral blood mononuclear cells (PBMCs) were isolated through gradient centrifugation using Lymphoprep (STEMCELL Technologies) within 4 hours. PBMCs were then resuspended in FBS/10% DMSO solution with a concentration of 7-10x10^6^ cells/mL for freezing in liquid nitrogen.

Frozen PBMCs were thawed in AIM-V media (Gibco, Thermo Scientific, MA, USA) supplemented with 10% FBS (Cytiva, USA) and DNase I (Stemcell Technology, Canada) (1 μg/mL) solution. 10^5^ PBMCs were allowed to rest in 96-round bottom well-plate containing AIM V media supplemented with 10% FBS, 10 mM HEPES, and 50 μM β-mercaptoethanol, for overnight before stimulation with synthesized long peptides at a concentration of 5 μM in a humidified incubator at 37°C with 5% CO_2_. PBMCs were further stimulated with GM-CSF (2000 IU/mL, Gibco, MT, USA) and IL-4 (1000 IU/mL, Invitrogen, MA, USA) for 24 hours. Following this initial stimulation, LPS (100 ng/mL, Sigma-Aldrich, MA, USA) and IFN-y (10 ng/mL, Gibco, MT, USA) were added to the PBMCs along with the peptides for an additional 12-hour. On the following day, IL-7, IL-15, and IL-21 (each at a concentration of 10 ng/mL) (Peprotech, NJ, USA) were added to the PBMC culture. The restimulation process involved exposing the peptides to a fresh media containing IL-7, IL-15, and IL-21 every 3 days for a total of 3 times. On day 12, PBMCs were restimulated with peptides and cultured in media without cytokines. ELISpot assays were performed on stimulated PBMCs on day 13.

### ELISpot assay on PBMCs stimulated with long peptides

Cultured T cells were transferred to an ELISpot plate (Mabtech, Sweden) and incubated for 20 hours at 37°C. PBMCs cultured with DMSO were used as a negative control group, while PBMCs stimulated with anti-CD3 were used as a positive control group. ELISpot assay was performed on treated PBMCs using ELISpot Pro: Human IFN-γ (ALP) kit (Mabtech, Sweden), following manufacture’s protocol. Developed spots on the ELISpot plate were then enumerated using an ELISpot reader (Mabtech, Sweden). The reactivity was determined by measuring the fold increase in the number of spots of PBMCs treated with mutant peptides relative to those treated with wild type peptides. A fold change of two was selected as the cut off for positivity (52).

### Flow cytometry intracellular staining for IFN-γ

Cells from ELISpot plate were collected in media supplemented with GolgiStop Protein Transport Inhibitor (BD Biosciences, NJ, USA) and incubated for 6 hr at 37°C. Positive control group was treated with 50 μM PMA (Abcam, UK), 1 mg/mL Ionomycin (Abcam, UK). Cells were then washed, blocked with Fc receptor (Biolegend, CA, USA), and stained with CD3-PE (clone HIT3a, Biolegend), CD4-PE/Cyanine7 (clone RPA-T4, Biolegend), CD8-FITC (clone RPA-T8, Cell Signaling) antibodies for 2 hr at 4°C. Cells were permeabilized for 20 mins at 4°C and then stained overnight with IFN-γ-APC (clone 4S.B3, Biolegend) antibody at 4°C.

### Statistical analysis

The Wilcoxon rank-sum test was used to compare the coverage, VAF, and immunogenicity percentile among three groups for Figure 3A, 3B and 4C. All statistical analyses were carried out using R (v2.6.3).

## RESULTS

### Comparison of mutation profiles from DNA sequencing and RNA sequencing data

RNA sequencing (RNAseq) data, which are commonly used for analysis of mutated gene expression in the current standard workflow of neoantigen identification, have been exploited to identify cancer-specific mutations in recent studies (26, 53, 54). However, the properties of RNAseq derived variants and neoantigens have not been fully characterized. To assess the utility of RNAseq in calling cancer-specific somatic mutations for neoantigen prediction, we sought to compare the mutation profiles obtained from RNAseq and DNAseq data across 25 CRC patients (**Table S1**), with a focus on all single nucleotide variants (SNVs) and indel variants (**Figure 1**). To achieve a balance between cost and mutation detection efficiency, we used a targeted sequencing panel consisting of 95 commonly mutated cancer-associated genes (**Table S2**). As a result, our comparison of RNAseq and DNAseq analysis was limited to these genes (**Figure 1**). The DNAseq and RNAseq data obtained from all 25 CRC patients have successfully met quality metrics, ensuring reliable datasets for mutation calling (**Table S3** & **S4)**. To identify mutations in DNAseq data, we used Dragen as our primary tool due to its superior performance in both SNV and indel mutation calling from a reference sample compared to other tools used in the analysis of DNAseq data (**Figure S1A**) (55).

**Figure 1.**
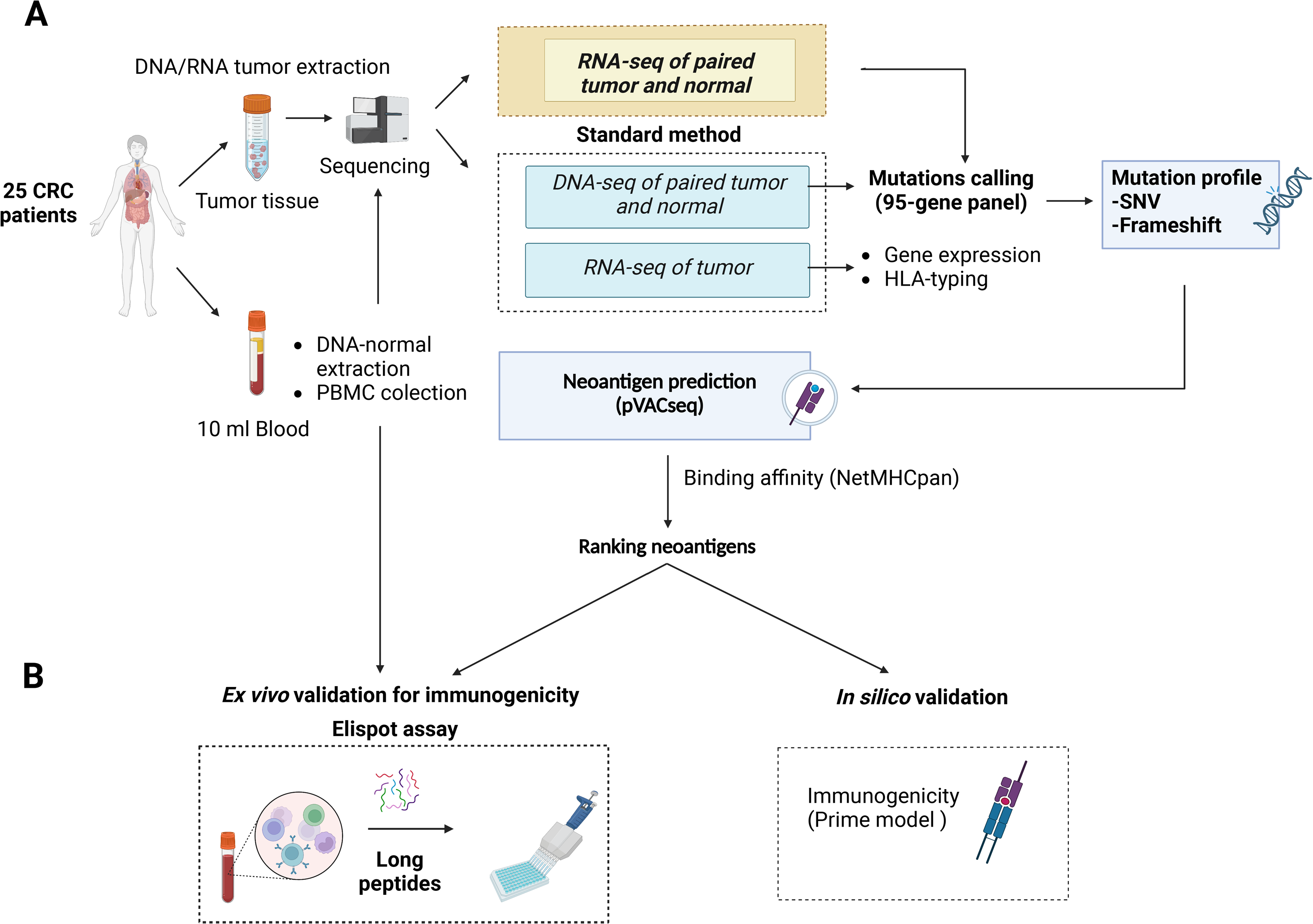
A novel workflow for CRC neoantigen identification and validation that integrates RNAseq data into somatic mutation calling. (A). Schematic diagram of the new workflow. Tumor biopsies and blood samples from CRC patients are subjected to targeted DNA and RNA sequencing, which focuses on a panel of 95 genes, for somatic mutation calling. Additionally, RNAseq data is used to determine gene expression and HLA-typing information. pVAC-Seq tool is then utilized for neoantigen prediction using DNA and RNA-derived somatic mutation data, gene expression data, and patient-specific HLA-typing data as inputs. (B). Methods to validate the advantages of the workflow. Predicted neoantigens from the workflow are subsequently validated by *ex vivo* ELISpot assay measuring IFN-γ secretion from PBMCs stimulated with long peptides carrying predicted variants and by *in silico* prediction of immunogenicity by PRIME tool.

To determine the most effective tool for calling mutations from RNAseq data, we compared the performance of VarScan and MuTect2. We found that VarScan yielded a higher proportion of variants overlapping with mutations detected from DNAseq compared to MuTect2 (18.3% versus 0.8%, **Figure S1B**). Furthermore, while MuTect2 tended to call a high percentage of indels with abnormal length, VarScan yielded a higher proportion of SNVs that were comparable to the mutation profiles identified from DNAseq (**Figure S1C** & **S1D**). These data suggested that VarScan exhibited higher sensitivity in detecting SNVs and produced fewer artifact indels. Thus, we decided to use VarScan as the calling tool for RNAseq data from the 25 CRC patients.

Out of the total 1,520 variants identified, only 340 (22.4%) were common between the two mutation calling methods, while most variants (77.6%) were exclusively detected by either DNAseq (DNA-unique) or RNAseq (RNA-unique) data. DNA-unique variants were more frequent than RNA-unique variants (56.1% versus 21.5.%, **Figure 2A**). Shared variants were detected in 16 out of the 25 CRC patients, accounting for 1% to 47% of the total identified variants (**Figure 2B** and **Table S5**). Interestingly, we found that RNA-unique variants were the major source of variants in 4 out of 25 (16%) patients (**Figure 2B**), while DNA-unique variants were identified as the major source of variants in the remaining 21 patients.

**Figure 2.**
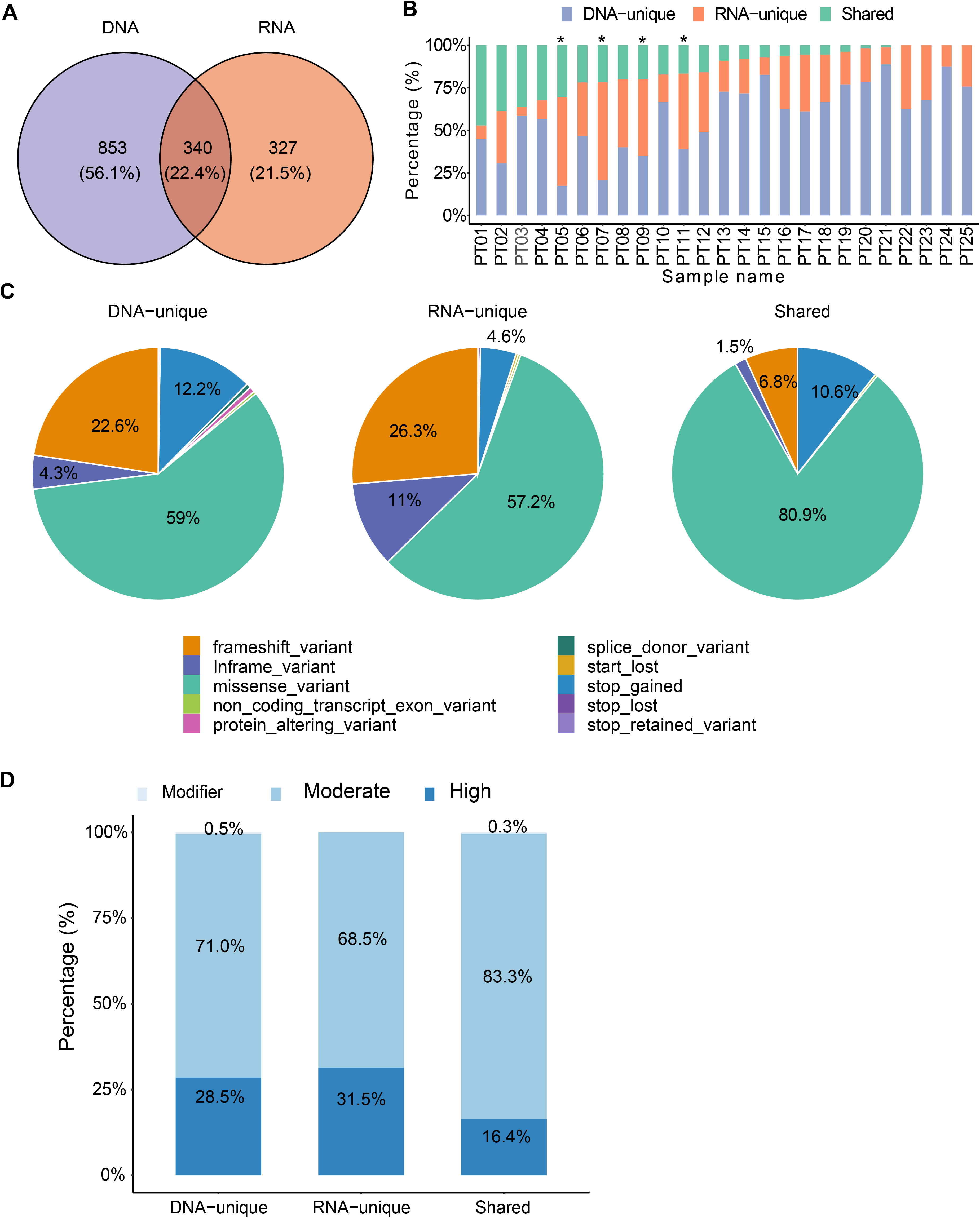
Comparison of identified somatic mutations between DNAseq data and RNAseq data. (A). Venn diagrams display the numbers of DNA and RNA mutations called by the specified mutation callers on matched tumor-normal DNAseq and RNAseq data from 25 CRC patients. (B). Proportions of each type of variants identified from both DNAseq and RNAseq data for each patient. Note: the graph is presented in descending order based on the proportion of shared variants. Patients marked with an asterisk exhibited a higher proportion of RNA-unique variants compared to DNA-unique variants. (C). Pie charts presenting the percentages of mutation types. (D). The proportions of indicated types of variants in relation to their phenotypic impacts.

When comparing the distribution of variant types between DNAseq and RNAseq, we observed a consistent pattern where missense variants were the most prevalent variant type (>50% of all variants in each group, **Figure 2C**). However, we did notice some notable differences. Explicitly, RNA-unique variants exhibited a higher frequency of in-frame variants (11% compared to 4.3% in DNAseq, **Figure 2C**) and frameshift variants (26.3% versus 22.6%, **Figure 2C**). On the other hand, DNA-unique variants had a higher occurrence of stop-gained variants (12.2% versus 4.6%, **Figure 2C**). In the shared-variant group, most variants consisted of missense variants (80.9%) and stop-gained variants (10.6%), collectively accounting for approximately 91.5% of all variants. To predict the functional impact of the three variant groups, we employed the Ensembl’s Variant Effect Predictor tool (36). Our analysis revealed that the phenotypic outcome was most significantly affected by RNA-unique variants in the high impact category, followed by DNA-unique and shared variants (**Figure 2D**). These results indicate a clear distinction between the tumor variant landscapes profiled by RNAseq and DNAseq, wherein RNAseq reveals a greater proportion of clinically relevant variants compared to DNAseq. Therefore, RNAseq appears to be particularly valuable in identifying variants with potential clinical significance.

To gain deeper insights into the variants identified by both sequencing methods, we conducted an analysis of their depth coverage and mutation allele frequency (MAF). Despite having lower coverage levels (P= 9.1x10^-5^, **Figure 3A**), the shared variants exhibited significantly higher MAFs (P= 2.22x10^-16^, **Figure 3B**) compared to the DNA-unique. This observation suggests that the shared variants are likely derived from major clones of somatic mutation clones, while the DNA-unique variants, characterized by significantly lower MAF (P < 2.22x10^-16^, **Figure 3B**), may originate from minor tumor clones.

RNA-unique variants displayed a notably lower median depth of coverage (P<2.22x10^-^16, **Figure 3A**) and MAF (20% versus 40%, p<2.22x10^-6^, **Figure 3B**) compared to the shared variants. These findings suggest that RNA-unique variants may originate from genes with low expression levels, resulting in a smaller number of variant transcripts. Intriguingly, the majority of shared variants and RNA-unique variants were identified in genes with high expression levels (FPKM >5, dashed line, **Figure 3C**), while unique variants identified through DNAseq (494/853, 58%, **Table S5**) were more commonly found in genes with low expression levels (FPKM <5, **Figure 3C**). Furthermore, when examining the MAF of variants in relation to their gene expression levels, shared variants (green dots, **Figure 3D**) exhibited higher levels of gene expression (FPKM >5) and MAF (> 24%) compared to other mutation types. In contrast, RNA-unique variants (orange dots, **Figure 3D**) tended to have similar gene expression levels but lower MAF, while a substantial number of DNA-unique variants (purple dots, **Figure 3D**) displayed both low gene expression and MAF. These observations strongly suggest that the MAF and transcriptional activity of mutated genes are significant factors contributing to the disparities observed between RNA-seq and DNA-seq. Notably, shared variants with high numbers of MAF may arise from dominant tumor clones and are highly expressed, making them potential neoantigen candidates. On the other hand, unique variants displaying low MAFs may be derived from subclonal mutations or poorly expressed mutations, further emphasizing the influence of MAF and gene expression on the distinct characteristics of the identified variants.

### *In silico* analysis of HLA-I binding affinity and immunogenicity of neoantigens derived from DNAseq and RNAseq

To identify neoantigen candidates, we utilized the pVAC-Seq pipeline, a well-established computation tool, to predict the binding affinity of 8-13 mer peptides generated from DNA or RNA variants to patient-specific HLA class I molecules (41). The HLA-I allele profiles of 25 patients were presented in **Table S6**. Through our analysis, we identified a total of 48,155 DNA-unique variants derived neoantigens (61.7%), 15,584 shared-variant derived neoantigens (20%), and 14,532 RNA-unique derived neoantigens (18.4%) (**Figure 4A, Table S7**). Remarkably, the proportions of neoantigens from each group showed a significant correlation with the proportions of nucleotide mutations (**Figure S2A**).

**Figure 3.**
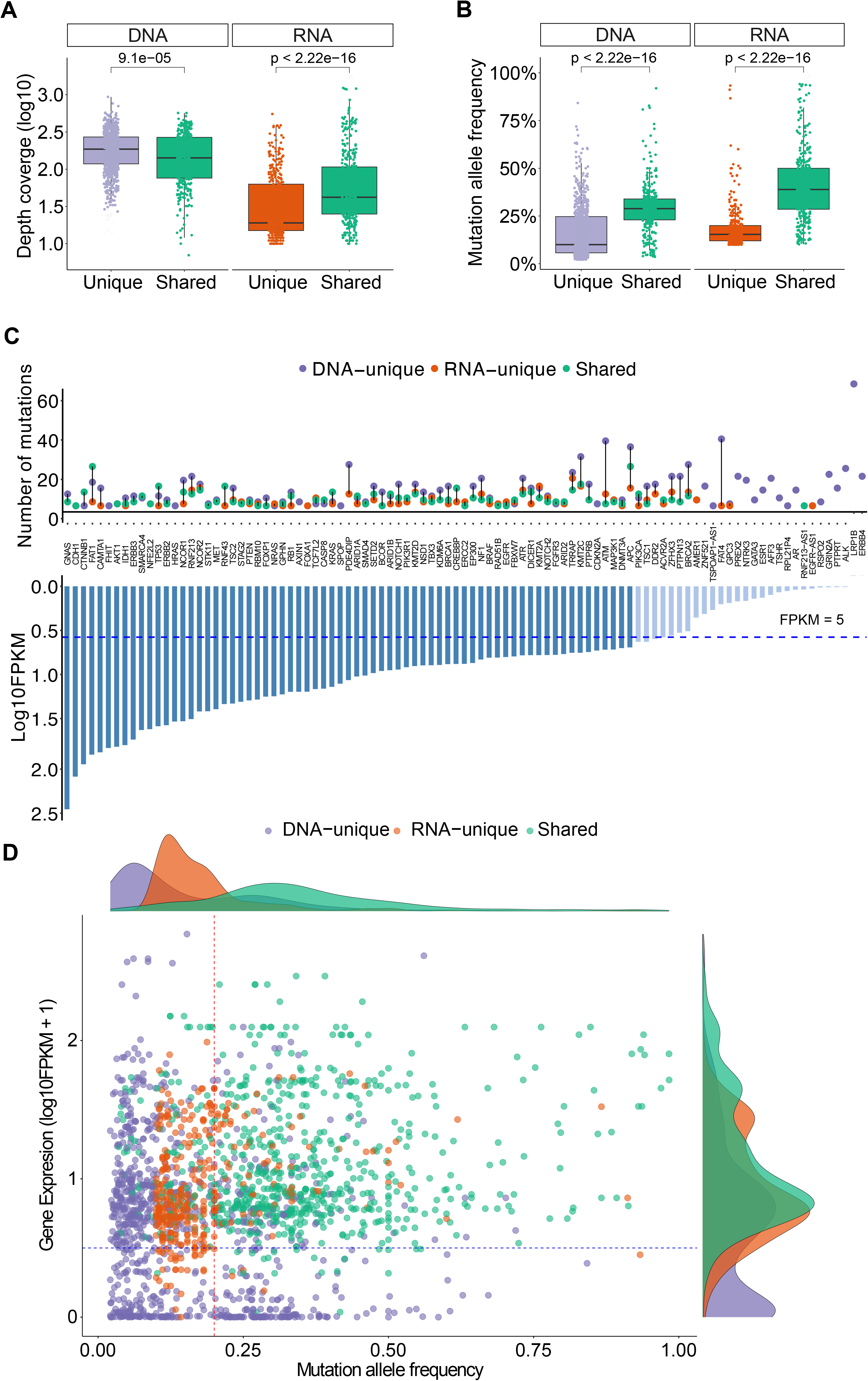
Depth coverage, MAFs and gene expression levels of variants from DNAseq and RNAseq data. (A). Depth coverage of the indicated groups of variants based on DNAseq and RNAseq data. (B). Mutation allele frequency of the indicated groups of variants. (C). A list of genes with indicated variants, along with their corresponding FPKM. (D). Gene expression levels of different groups of variants in relation to their mutation allele frequency. In panels (A) and (B), the boxes represent the median value, as well as the lower and upper quartiles (25th and 75th percentiles). The p-values were obtained from the Wilcoxon rank-sum test.

**Figure 4.**
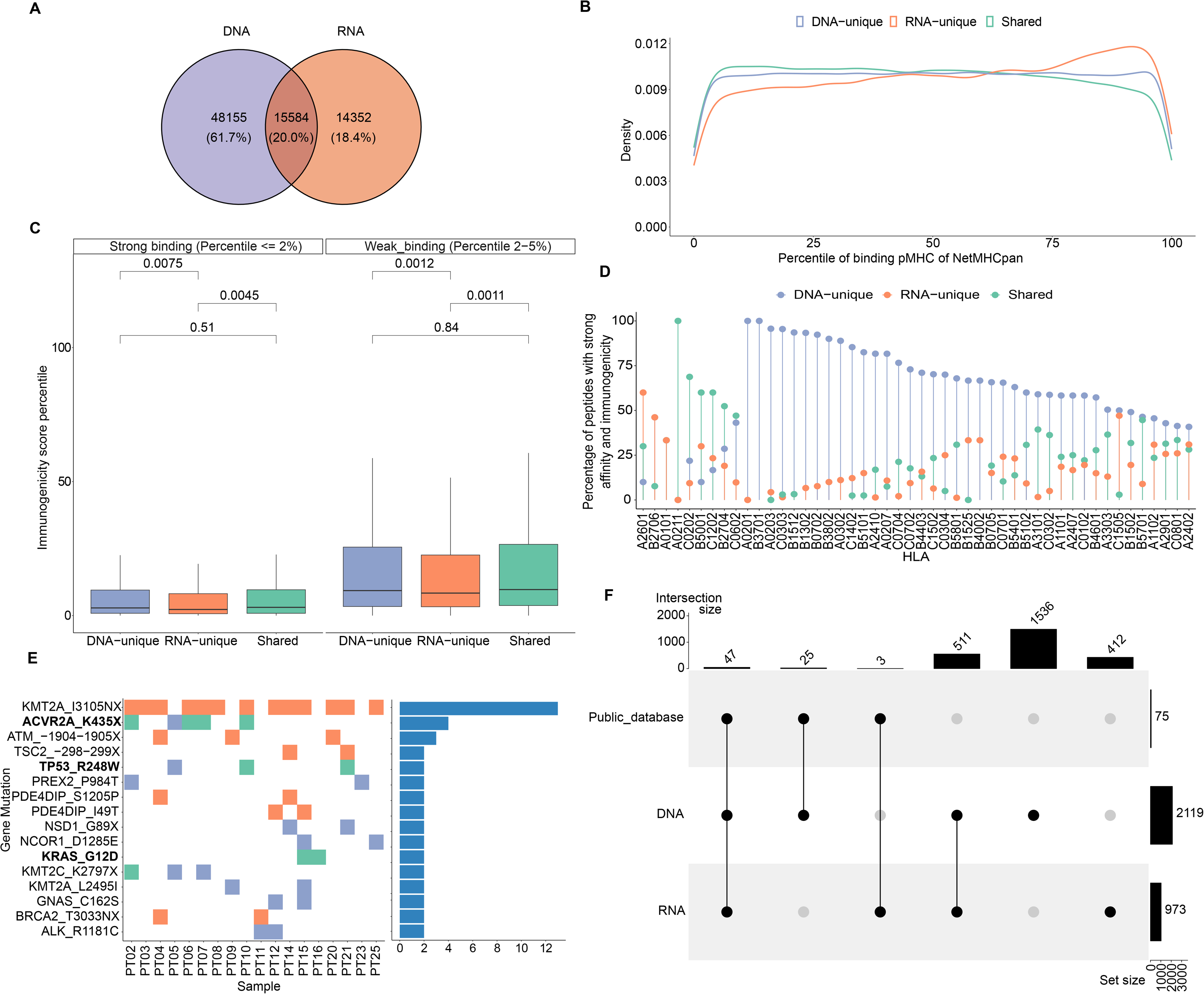
HLA-I binding affinity and immunogenicity of predicted neoantigens derived from DNA-seq and RNA-seq data. (A). A Venn diagram illustrates the proportion of each type of neoantigens identified from DNAseq and RNAseq data. (B). Histograms showing the density of neoantigens according to their percentile ranks of HLA-I binding affinity calculated by NetMHCpan. (C). Predicted immunogenicity, as calculated by the PRIME tool, for both strong binding and weak binding neoantigens. The box plot represents the median value, along with the lower and upper quartiles (25th and 75th percentiles). Outliers are not displayed for clarity of visualization. The p-values were estimated using the Wilcoxon rank-sum test. (D). A Lollipop plot depicts the distribution of specific groups of neoantigens based on their percentage, focusing on indicated HLA-I alleles. These plots highlight neoantigens that fall within the top 2% in terms of strong binding affinity to HLA-I and demonstrate high immunogenicity. (E). A map illustrates the frequency of indicated mutations on 25 CRC patients. The ones highlighted in bold have been previously validated as highly immunogenic through immunological assays in previous studies. (F). An UpSet plot illustrates the frequency distribution of the indicated groups of variants identified from public datasets.

It is well established that effective activation of T cell responses relies on the presentation of neoantigens on the patient’s HLA-I molecules (56). Here, the binding affinity of predicted neoantigens from each group of tumor variants to HLA-I was predicted by NetMHCpan 4.1 (17). In **Figure 4B**, we showed the density of predicted neoantigens derived from DNA-unique, RNA-unique or shared variants according to the percentile rank of HLA-I binding affinity predicted by NetMHCpan 4.1. In the 0 to 50 percentile windows, neoantigens from shared variants had the highest proportion of neoantigens fallen in this range, followed by those from DNA-unique, then RNA-unique variants (**Figure 4B**). Conversely, the RNA-unique derived neoantigens were the most highly enriched one in the 50 to 100 percentile range (**Figure 4B**). Thus, compared to neoantigens derived from DNA-unique variants, RNA-unique derived neoantigens exhibited a lower HLA-I binding affinity according to NetMHCpan. It has been reported that the binding affinity to HLA-I is determined by specific anchor residues in neopeptides (57). When comparing DNA-unique and shared neoantigens with RNA-unique neoantigens, it was observed that the latter exhibited a reduced proportion of mutations at P2 (**Figure S2B**). Notably, P2 serves as a crucial anchor residue involved in the primary interactions between the peptide and HLA-I molecule, and mutations occured within this position increase the binding affinity to HLA-I. This observation suggests that the decreased frequency of RNA-unique derived neoantigens carrying mutations at this anchor site, in comparison to other sources of neoantigens, may account for their lower binding affinity.

To assess the immunogenicity of the predicted neoantigens, we employed the PRIME tool which captures biophysical properties of both antigen presentation and TCR recognition to evaluate their potential to elicit a CD8^+^ T cell-specific immune response (42). The predicted immunogenicity of neoantigens was evaluated in relation to their predicted binding affinity to HLA-I (**Figure 4C**). Neoantigens were categorized into two groups based on the standard threshold of NetMHCpan: strong binding, which included neoantigens with predicted binding affinity within the top two percentile, and weak binding, which encompassed neoantigens with binding affinity higher than the two-percentile threshold. We observed a positive correlation between the predicted binding affinity to HLA-I using NetMHCpan and the predicted immunogenicity assessed by the PRIME tool, irrespective of the neoantigen class. Notably, strong binding neoantigens exhibited lower percentile ranks of immunogenicity (**Figure 4C**). However, intriguingly, among the neoantigens with strong HLA-I binding affinity, the RNA-unique neoantigens showed significantly lower percentiles of immunogenicity compared to both DNA-unique (P=0.0075, **Figure 4C**) and shared neoantigens (P= 0.0045, **Figure 4C**). Within the weak binding neoantigens, RNA-unique neoantigens consistently demonstrated lower percentiles of immunogenicity compared to DNA-unique (P=0.0012, **Figure 4C**) and shared neoantigens (P=0.0011, **Figure 4C**). Subsequently, neoantigens meeting the criteria of predicted binding affinity and immunogenicity within the top two percentile for both parameters were profiled based on the specific HLA-I alleles identified in our cohort of 25 CRC patients. As shown in **Figure 4D**, we observed that the binding affinity of predicted neoantigens to HLA-I was influenced by both the specific neoantigen sequence and the HLA-I allele. For instance, we observed that the HLA-I allele A02011 exhibited a higher binding affinity to shared neoantigens, as this allele showed the highest proportion of detected neoantigens in this group. Similarly, the HLA-I allele A2601 displayed a stronger binding affinity for RNA-unique derived neoantigens; while the HLA-I allele A0201 showed a stronger binding affinity for DNA-unique derived neoantigens, in comparison to shared and RNA-unique neoantigens (**Figure 4D**). Among the neoantigens displaying strong predicted affinity and immunogenicity, a noteworthy subset of 16 neoantigens was consistently identified in at least two patients (**Figure 4E**). Of those, neoantigens derived from three shared mutations (ACVR2A_K435X, TP53_R428W, and KRAS_G12D) have been experimentally validated in previous studies and reported in public databases of immunogenic neoantigens. Notably, the KMT2A_IN3105X neoantigen predicted from an RNA-unique variant, exhibited the highest frequency among these frequently detected neoantigens, being present in 13 out of 25 (52%) patients. This suggests that this neoantigens has the potential to serve as a public neoantigen, capable of eliciting immune responses across multiple individuals. Additionally, a total of 75 strong affinity and immunogenic neoantigens were previously reported in public databases of immunogenic peptides. Among these, the majority (47/75, 62.7%%) could be found from shared variants, while 25 and 3 neoantigens were predicted from DNA-unique and RNA-unique variants, respectively (**Figure 4F**). These findings underscore the presence of both shared and unique neoantigens with strong binding and immunogenicity in 25 analyzed patients, further highlighting the importance of considering different sources of NGS data for mutation identification in neoantigen-based immunotherapy approaches.

Taken together, these findings emphasize the distinct binding affinity and immunogenic potential of neoantigens originating from different variant groups. Particularly, our data suggests that despite their low predicted binding affinity, neoantigens derived from RNA somatic mutations still exhibit high immunogenicity, indicating their potential to elicit an immune response for immunotherapy. These observations underscore the importance of considering not only DNA-seq but also RNA-seq derived variants for selecting candidate neoantigens.

### Experimental validation of predicted neoantigens by ELISpot

To evaluate the effectiveness of integrating RNA-seq variant calling into the current standard method, we conducted ELISpot assay on four CRC patients using autologous PBMCs following the procedure outlined in **Figure 5A**. Initially, we identified 431 nonsynonymous variants from both DNAseq and RNAseq data, resulting in a total of 18,479 predicted neoantigens using the pVAC-Seq tool. To accommodate the limited availability of PBMCs, only the top ten mutations resulting in neoantigens with the highest predicted binding affinity to HLA-I were chosen for each patient. As a result, a total of 40 synthesized long peptides (LPs) carrying the corresponding mutations were synthesized and used in an *ex vivo* ELISpot assay to measure the release of IFN-γ from patients’ PBMCs (**Figure 5A, Table S8**).

**Figure 5.**
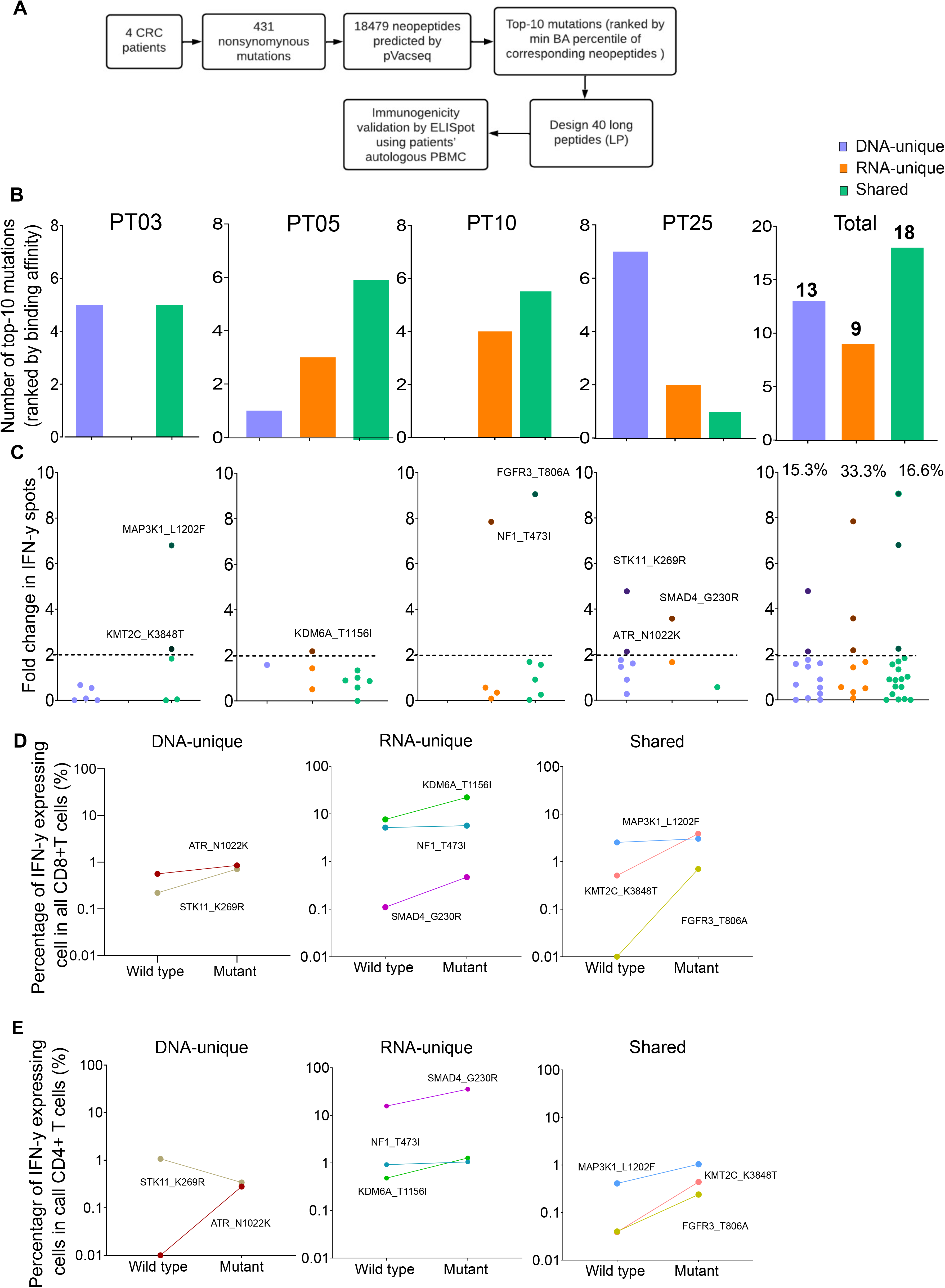
Validation of neoantigens *in silico* identified from the modified workflow by ELISpot assays on four CRC patients. (A). A schematic diagram illustrates the procedural steps of neoantigen prioritization and the ELISpot assay. (B). The number of each type of neoantigens identified from each CRC patient. (C). The fold change in IFN-γ spots, relative to the wildtype peptides, for 40 long peptides. Note: only the mutants that result in a positive value in ELISpot are depicted with their corresponding amino acid change. (D). The percentage of IFN-γ expressing CD4+ T cells induced by indicated long peptides. Note: these long peptides induce a more than 2-fold change in IFN-γ spots as observed in the ELISpot assay. (E). The percentage of IFN-γ expressing CD8+ T cells induced by indicated long peptides.

Among the 40 designed LPs, those originating from shared neoantigens were detected in all patients, whereas LPs derived from DNA-unique or RNA-unique variants were only detected in three out of four patients (**Figure 5B**). However, no LPs were identified within the DNA-unique group for patient PT10 and within the RNA-unique group for patient PT03 (**Figure 5B**). When considering the cumulative number of LPs across all patients, it was observed that shared-variants yielded the highest number (18 out of 40), while RNA-unique variants yielded the fewest (9 out of 40, **Figure 5B**).

The PBMCs from four patients were subjected to three rounds of stimulation with 40 LPs carrying mutations or their corresponding wildtype counterparts to measure the secretion of IFN-γ. The ELISpot results for the 40 tested LPs were presented in **Figure 5C** and **Table S8**. A fold change of two in the number of IFN-γ spots from LPs relative to their corresponding wildtype peptides was chosen as the positivity cutoff, with LPs resulting in an ELISpot fold change value of two or higher considered as immunogenic (52). Among 40 tested LPs, we identified eight immunogenic LPs, with three originating from RNA-unique variants, three from shared variants, and two from DNA-unique variants (**Figure 5C** & **Figure S3**). Notably, all four patients had at least one LP capable of inducing IFN-γ production by PBMCs. Among the LPs derived from RNA-unique variants, three out of nine (33.3%) were positive for IFN-γ activation, while the proportions of positive LPs were lower for those derived from shared variants (three out of 18, 16.7%, **Figure 5C**) or DNA-unique variants (two out of 13, 15.4%, **Figure 5C**). The findings suggest that RNA-unique variants may result in fewer neoantigens with strong binding affinity to HLA-I, but they are more likely to activate T cells compared to shared or DNA-unique neoantigens.

Intracellular flow cytometry staining of IFN-γ in T cells further demonstrated that all LPs showing positive results in the ELISpot assay effectively activated CD8+ T cells. This activation led to a significant increase in the percentage of IFN-γ positive cells, with a fold increase greater than 1 compared to their corresponding wildtype peptides (**Figure 5D** & **Figure S4**). Moreover, consistent with the activation of CD8^+^ T cells, all LPs exhibited increased production of IFN-γ by CD4^+^T cells, except for the LP carrying STK11_K269R, which originated from a DNA-unique variant (**Figure 5E**). Although this LP did not exhibit detectable changes in intracellular IFN-γ levels in CD4^+^ T cells, it still demonstrated CD8^+^ T cell activation. Overall, these findings suggested that the integration of RNAseq data for variant calling into the current neoantigen prediction workflow could enhance the identification of effective and immunogenic neoantigen candidates for the development of cancer immunotherapies.

## DISCUSSION

The identification of highly immunogenic neoantigens capable of eliciting T-cell-mediated responses is essential for the development of effective personalized immunotherapies for cancer. However, the current challenge lies in accurately identifying these neoantigens due to the limited number of highly immunogenic neopeptides predicted by conventional bioinformatic workflows. These workflows solely rely on genomic sequencing data for tumor mutation calling, overlooking the potential contribution of transcriptomic variants in generating neoantigens. To address this limitation, we aimed to enhance the identification of highly immunogenic neoantigens by integrating RNA sequencing data into the conventional bioinformatic workflow (**Figure 1**). By considering tumor mutations at the transcriptional level, we sought to expand the pool of valuable immunogenic neopeptides for colorectal cancer (CRC) patients. In our study, we successfully demonstrated that integrating into the conventional workflow for variant calling significantly increased the number of valuable immunogenic neopeptides for CRC patients. This improvement provides a promising avenue for the development of more effective cancer treatments.

Our analysis of tumor variants using DNAseq and RNAseq data obtained from 25 CRC patients identified a moderate proportion (22.4%) of shared somatic variants (**Figure 2A**). This finding is consistent with a previous study that reported a similar trend in two datasets (58). Additionally, the proportions of shared mutations exhibited significant variation among patients (**Figure 2B**), highlighting the intrinsic diversity of cancer mutations and the heterogeneity of clonal expansion within each patient. Furthermore, different variant groups displayed distinct characteristics, with RNA-variants showing an enrichment for frameshift and inframe variants and displaying more profound impact on the phenotypic outcome (**Figure 2C** & **D**). Neoantigens derived from frameshift or indel variants, which are greatly distinct from self peptides, have been shown to generate highly immunogenic tumor neoantigens and thereby expand the pool of ideal candidates for immunotherapy (59, 60).

Both DNA-unique and RNA-unique variants displayed significantly lower MAFs compared to shared variants (**Figure 3B**). This observation implies that these unique variants likely originated from tumor clones with low frequencies, which might not be consistently detected at both genomic and transcriptomic levels due to the limited sensitivity of sequencing methods. Notably, our analysis revealed that DNA-unique variants were more frequently associated with genes characterized by low FPKMs, unlike shared or RNA-unique variants (**Figure 3C** & **D**). These findings suggest that DNA-unique variants may arise from genes with low expression or those displaying mono-allelic expression of the wild-type allele. Conversely, RNA-unique or shared variants tend to occur in genes exhibiting high expression levels, implying their abundant transcription. A previous study have demonstrated a correlation between the expression levels of neoantigens and their likelihood of being presented by HLA-I on the surface of tumor cells, which can trigger immune responses leading to the eradication of tumor cells (61). Hence, neoantigens arising from RNA-unique or shared variants might be superior, as they are more likely to be presented and recognized by the immune system. The discrepancies in mutation profiles between RNA-seq and DNA-seq could be attributed to the low MAFs, low quantities of transcripts harboring variants and/or insufficient sequencing coverage.

The proportions of neoantigens predicted by the pVAC-Seq tool are similar to those of nucleotide variants (**Figure 2A** & **4A**). Currently, the prediction of peptide binding affinity for HLA-I is a pivotal criterion in the selection of neoantigens for experimental validation (17). Employing NetMHCpan 4.1, we discovered that neopeptides originating from RNA-unique variants exhibited lower percentile ranks of binding affinity compared to those derived from shared or DNA-unique variants (**Figure 4B**). This finding suggests that neopeptides resulting from RNA variants tend to display reduced levels of HLA-I binding affinity in comparison to those arising from DNA variants. Prior research has indicated that the position of mutations within mutant peptides can influence their binding affinity to HLA-I molecules, with specific residues in the peptides, known as anchor residues, serving as key determinants of binding affinity (62). Therefore, it is plausible that amino acid changes in neopeptides predicted from RNA mutations may arise from positions that do not lead to enhanced binding affinity, in contrast to those arising from DNA mutations. Interestingly, our findings revealed a lower proportion of RNA-derived neoantigens with mutations occurring at the primary anchor site P2, which is recognized as a critical factor influencing peptide affinity for various HLA-I types. This distinction was observed when comparing RNA-derived neoantigens with both shared and DNA-unique derived neoantigens **(Figure S2B**) (63). Another possible explanation for the lower binding affinity of RNA-unique neoantigens could be attributed to the fact that current prediction tools have not been specifically trained on this particular group of neoantigens (64).

While predicted HLA-I binding affinity serves as a crucial indicator for the presentation of neoantigens on tumor cells, it is not the sole determinant of neoantigen immunogenicity. The immunogenicity of neoantigens is also influenced by the interaction between peptide-HLA complexes and T cell receptors (TCR) (42, 65, 66). Therefore, in our study, we initially selected neopeptides with strong binding affinity (< 2% percentile rank). Subsequently, we employed the PRIME tool (42), which captures molecular properties related to both antigen presentation and TCR recognition, to estimate the immunogenicity of these selected peptides. Interestingly, we observed that neoantigens derived from RNA-unique mutations or shared mutations exhibited significantly higher immunogenicity compared to those derived from DNA-unique mutations (**Figure 4C**). Schmidt et al. have identified specific amino acid positions within the neopeptide sequence, known as minimally impacting on HLA-I affinity (MIA) positions. These positions have been found to have significant roles in binding to the T cell receptor (TCR) (42). Therefore, it is plausible that amino acid changes in neopeptides derived from RNA mutations may occur at such positions, resulting in enhanced TCR affinity and consequently explaining their stronger immunogenicity. Analysis of neoantigen immunogenicity, considering the HLA-I allele panels obtained from our CRC patient cohort, revealed a notable dependence on specific HLA-I alleles, thereby emphasizing the significance of profiling the HLA-I genotype of cancer patients for personalized immunotherapy (**Figure 4D**). The notable immunogenicity scores of neopeptides derived from RNA variants suggest their potential to effectively activate T cell-mediated immune responses, rendering them valuable candidates for clinical evaluation. Our *in silico* analysis successfully identified a recurrent RNA-derived neopeptide (KMT2A_IN3105X) in 25 CRC patients. Additionally, we discovered three shared neopeptides (ACVR2A_K435X, TP53_R428W, and KRAS_G12D) that have been experimentally validated as highly immunogenic in publicly available databases (**Figure 4E** & **F**). These neopeptides hold potential as public neoantigens, making them suitable candidates for an off-the-shelf vaccine strategy. Thus, we speculate that incorporating RNA-unique variants, which exhibit strong binding affinity and higher transcription abundance, can serve as a strategy to identify more effective targets for neoantigen-based vaccination.

To validate our hypothesis regarding the effectiveness of neoantigens derived from RNA variants compared to DNA-derived neoantigens, we conducted *ex vivo* ELISpot assays on four patients with available blood samples for PBMC collection. The purpose was to assess the immunogenicity of predicted neopeptides originating from different mutation sources. For each patient, we selected the top 10 mutations based on the predicted binding affinity of the corresponding neopeptides to the patients’ HLA-I profile. To evaluate immunogenicity, we designed LPs incorporating these mutations (**Figure 5A**). Consistent with our analysis on 25 CRC patients, the proportion of LPs derived from RNA-unique mutations with strong binding affinity was lower compared to those derived from DNA-unique or shared mutations (**Figure 5B**). However, in the *ex vivo* ELISpot assays, three out of nine LPs (33.3%) carrying RNA-unique variants triggered IFN-γ production in PBMCs of three out of four patients, while only two out of 13 LPs (15.3%) carrying DNA-unique variants induced IFN-γ production in a single patient (**Figure 5C**). In line with the ELISpot data, we detected IFN-γ activation not only in CD8^+^ T cells but also in CD4^+^ T cells for most of the tested long peptides. However, one LP derived from a DNA-unique mutation exclusively activated CD8^+^ T cells (**Figure 5D** & **E**). Our selection and design of LPs were based on the rank of neopeptides’ HLA-I binding affinity, aiming to specifically activate CD8^+^ T cells. However, our findings align with a previous study demonstrating that LPs covering target mutations could be intracellularly processed to peptides of differrent lengths and subsequenty presented to both CD4^+^ and CD8^+^ T cells (67). Our *ex vivo* validation of neoantigens’ immunogenicity using patients’ PBMCs provides compelling experimental evidence that relying solely on DNAseq data for tumor mutation calling would overlook valuable neoantigens derived from RNA variants and that integrating variant calling by RNAseq into this process significantly enhances the likelihood of detecting immunogenic neoantigens.

This study has several limitations that should be acknowledged. Firstly, in order to develop a cost-effective workflow for neoantigen identification, the analysis was focused on SNV and indel variants within only 95 cancer-associated genes. Consequently, other types of mutations, such as gene fusions and alternative splicing, and other genes were not explored (68, 69). Secondly, while RNAseq holds the potential to identify mutations on a genome-wide scale, its sensitivity and specificity are influenced by many factors such as sequencing depth, tumor purity, and the variant calling pipeline. To mitigate the potential impact of these biases, we carefully selected the optimal mutation caller for RNA-seq data, VarScan, after comparing its performance with MuTect2. However, more validation studies are necessary to improve the variant calling tools for RNA-seq data and standardize their use. Thirdly, the study was conducted with a limited sample size of 25 CRC patients, and the experimental validation of predicted neoantigens through *ex-vivo* ELISpot assays was performed on only four patients due to the availability of blood samples. As a result, the generalizability of the findings may be constrained. Finally, the assessment of the immunogenicity of candidate LPs relied exclusively on *ex-vivo* stimulation of patients’ PBMCs, which may not accurately reflect the natural presentation of neoantigens by HLA-I molecules expressed in patients’ tumor cells. Therefore, additional experimental validation using liquid chromatography mass spectrometry-based immunopeptidomics may be required to confirm the presentation of predicted neoantigens on HLA-I molecules in tumor cells.

Taken together, in this proof-of concept study, we provide compelling evidence for the benefits of utilizing RNAseq-guided mutations for neoantigen prediction, as it allows for the identification of a larger pool of potential and highly immunogenic neoantigens by leveraging additional information from RNAseq data beyond conventional gene expression levels.

### Author contributions

BQT.N and TPD.T conduct experiments, perform formal analysis, curate data, and develop methodologies. HT.N is responsible for patient recruitment and conceptualization. TN.N specializes in data curation and formal analysis. TMQ.P conducts experiments and performs formal analysis. HTP.N and V.N conduct experiments, perform formal analysis, and curate data. DH.T, TS.T, TN.P, and M.L recruit patients and analyze data. M.P, H.G, and HN.N conceptualize the study and edit writings. LS.T conceptualizes the study, writes the original manuscript, and edits the final document.

## Acknowledgments

The authors thank all participants who agreed to take part in this study. We thank Dr. Kien Nguyen for proofreading our manuscript.

## FIGURE LEGEND

**Figure S1. Evaluation of mutation calling tools for DNAseq and RNAseq data**

(A). Comparison of performance of three indicated mutation callers on a reference DNAseq dataset.

(B). A Venn diagram illustrates the number of mutations identified by Dragen and two RNA mutation callers, VarScan and MuTect2.

(C). Proportions of SNV and indel mutations called by indicated tools.

(D). Length distribution of Ins and Indel mutations called by indicated tools

**Figure S2. Distribution of mutation positions of DNAseq and RNAseq derived neoantigen**

(A). Correlation between the numbers of variants and neoantigens within the indicated groups.

(B). A lollipop plot displays the percentage of neoantigens from the indicated groups that contain mutations at positions 1 to 12. The blue box represents the anchor site of the peptide and HLA-I molecule.

**Figure S3. ELISpot assays on eight long peptides which result in 2-fold change of IFN-γ spots.**

**Figure S4. Gating strategy for detecting IFN-γ production from CD4^+^ and CD8^+^ T cells in LP-stimulated PBMCs of 4 CRC patients.**

**Table S1: Clinical Characteristics of 25 CRC patients.**

**Table S2: List of 95 genes used for mutation calling in the study.**

**Table S3: Quality control metrics of DNA sequencing**.

**Table S4: Quality metrics of RNA sequencing.**

**Table S5: List of somatic variants detected from DNAseq and RNAseq data in tumor tissues of 25 CRC patients**.

**Table S6: HLA-I allele profiles of 25 CRC patients**.

**Table S7: List of neoantigens with 2% percentile cut-off for binding affinity score (NetMHCpan)**.

**Table S8: ELISpot and intracellular staining of IFNγ from 40 long peptides harboring mutations selected from the top-10 neoantigens ranked by binding affinity to HLA-I**.

