## Supplementary figures and images for "Improvement in Neoantigen Prediction via Integration of RNA Sequencing Data for Variant Calling"

### Supplemental Figure 1

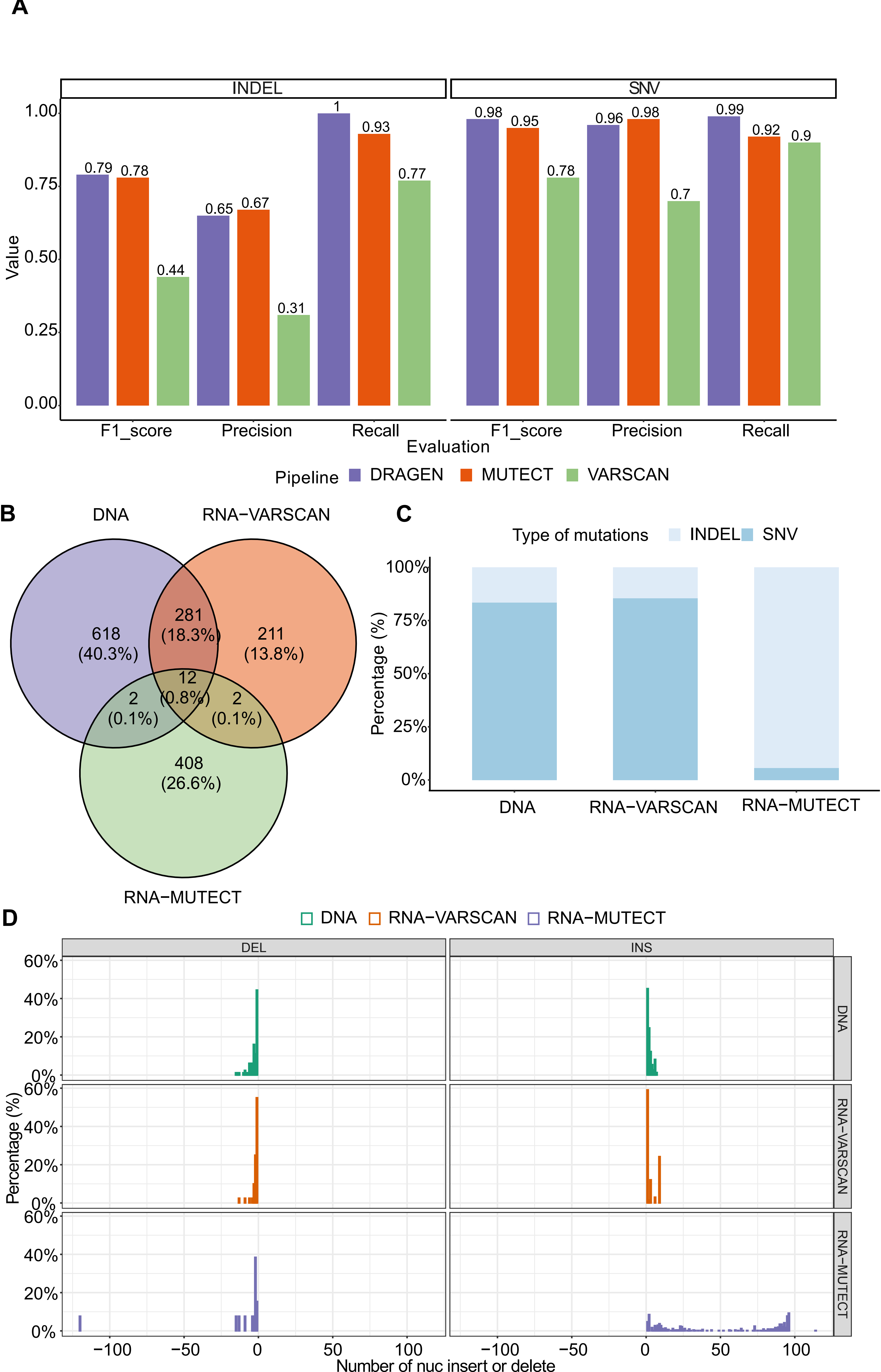

### Supplemental Figure 2

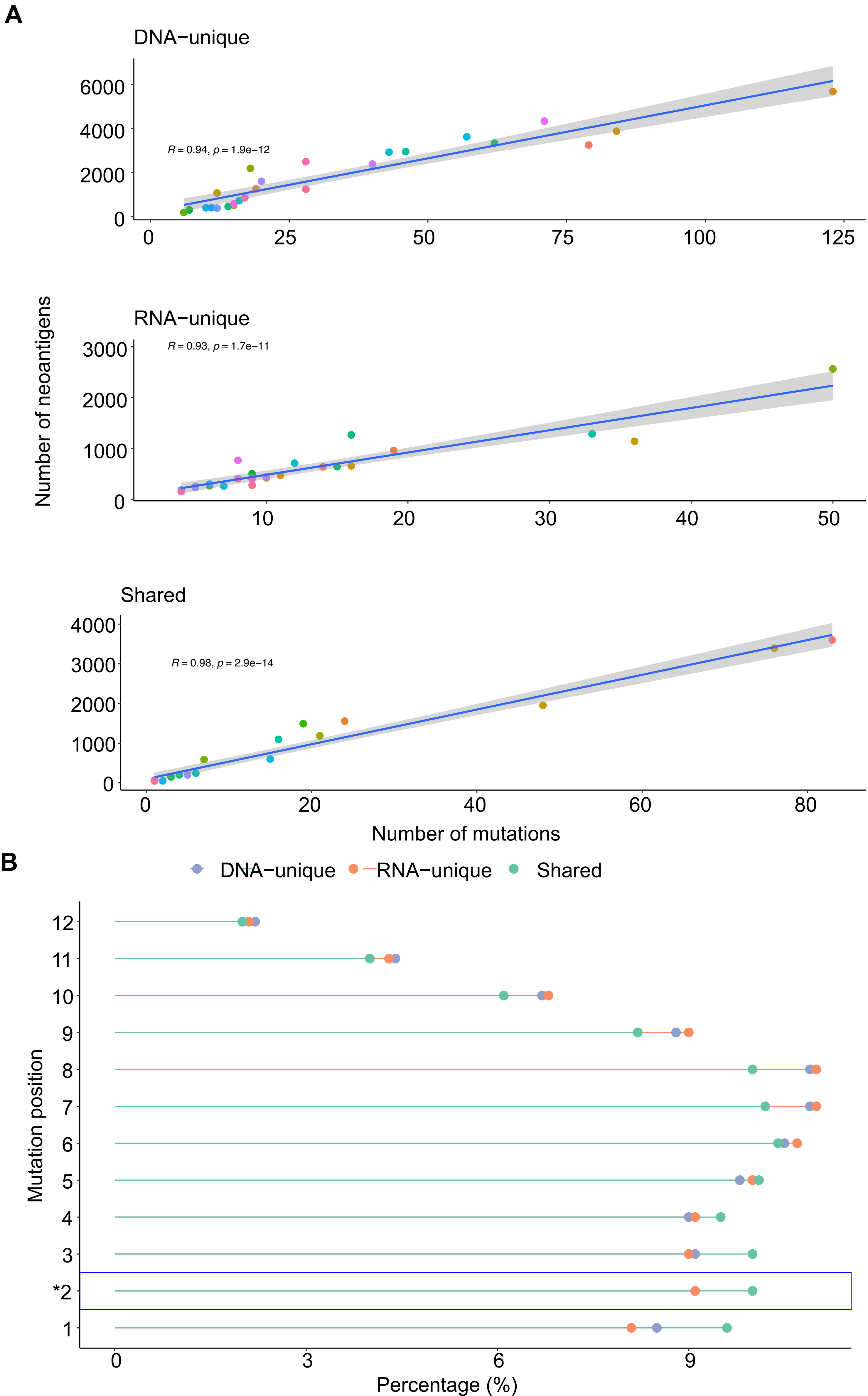

### Supplemental Figure 3

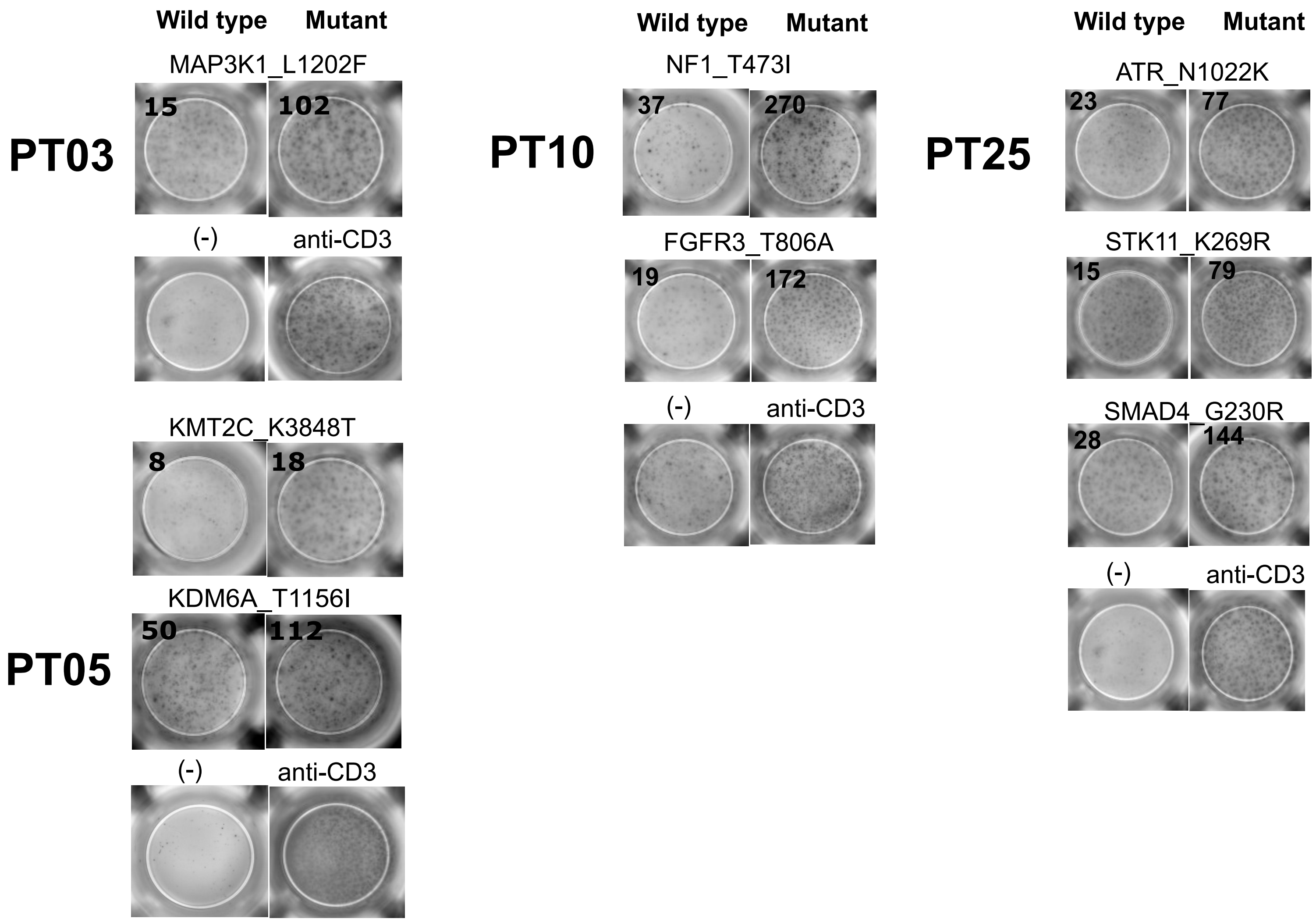

### Supplemental Figure 4

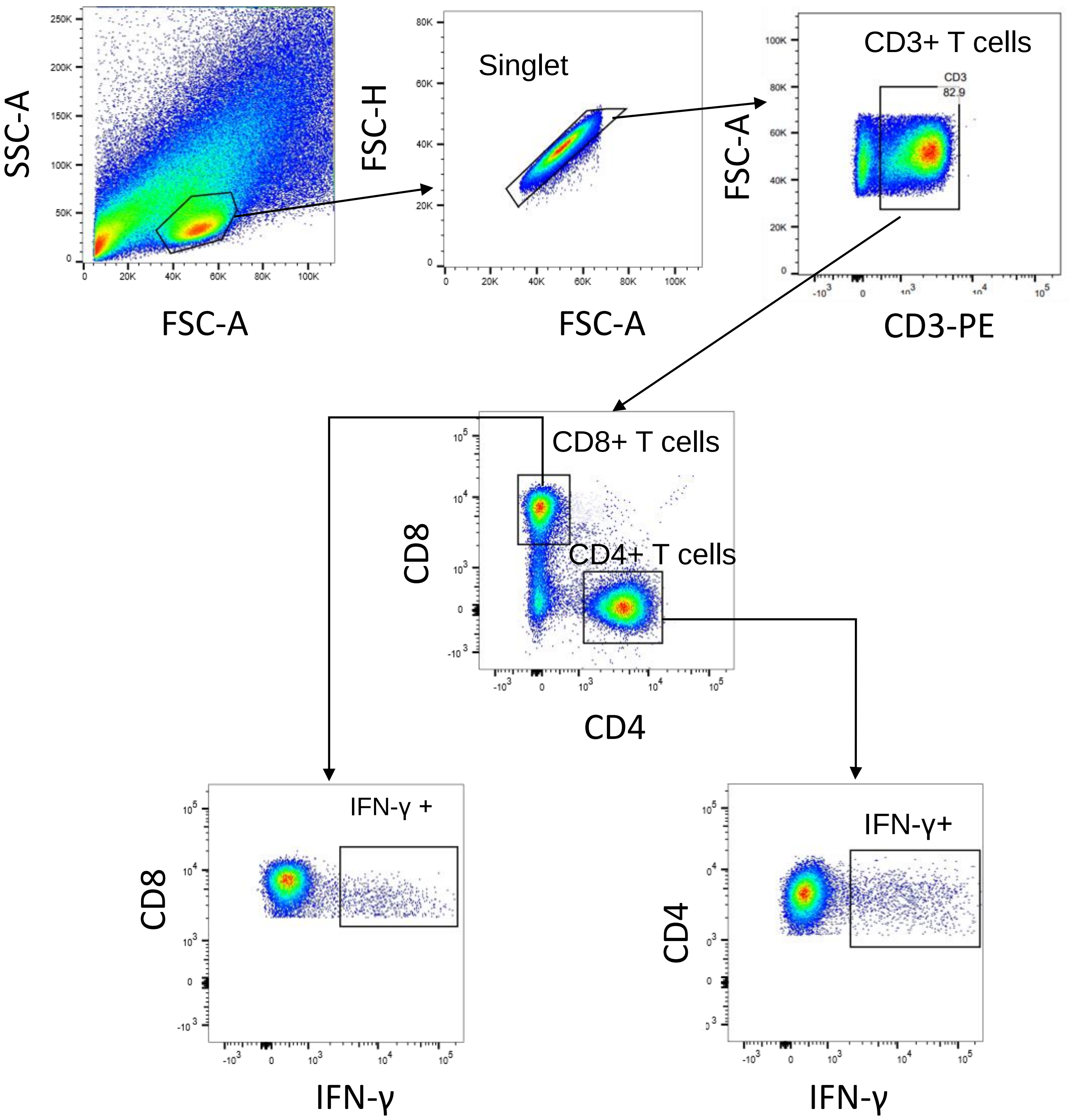
